# *Shake-it-off*: A simple ultrasonic cryo-EM specimen preparation device

**DOI:** 10.1101/632125

**Authors:** John L. Rubinstein, Hui Guo, Zev A. Ripstein, Ali Haydaroglu, Aaron Au, Christopher M. Yip, Justin M. Di Trani, Samir Benlekbir, Timothy Kwok

**Affiliations:** The Hospital for Sick Children Research Institute, Toronto, Ontario, Canada; Department of Medical Biophysics, The University of Toronto, Ontario, Canada; Department of Biochemistry, The University of Toronto, Ontario, Canada; Engineering Science Division, The University of Toronto, Ontario, Canada; Institute of Biomaterials and Biomedical Engineering, University of Toronto, Ontario Canada

**Keywords:** Cryo-EM, specimen preparation, ultrasonic, self-wicking grids, Raspberry Pi, 3D printing, CNC milling

## Abstract

Although microscopes and image analysis software for electron cryomicroscopy (cryo-EM) have improved dramatically in recent years, specimen preparation methods have lagged behind. Most strategies still rely on blotting microscope grids with paper to produce a thin film of solution suitable for vitrification. This approach loses more than 99.9% of the applied sample and requires several seconds, leading to problematic air-water interface interactions for macromolecules in the resulting thin film of solution and complicating time-resolved studies. Recently developed self-wicking EM grids allow use of small volumes of sample, with nanowires on the grid bars removing excess solution to produce a thin film within tens of milliseconds from sample application to freezing. Here we present a simple cryo-EM specimen preparation device that uses components from an ultrasonic humidifier to transfer protein solution onto a self-wicking EM grid. The device is controlled by a Raspberry Pi single board computer and all components are either widely available or can be manufactured by online services, allowing the device to be constructed in laboratories that specialize in cryo-EM, rather than instrument design. The simple open-source design permits straightforward customization of the instrument for specialized experiments.

**Synopsis:** A method is presented for high-speed low-volume cryo-EM specimen preparation with a device constructed from readily available components.

## Introduction

During the 1980s Jacques Dubochet and colleagues established that thin films of protein solutions can be vitrified and imaged by electron microscopy (Dubochet *et al.*, 1988). Dubochet’s method, which is still standard for electron cryomicroscopy (cryo-EM), requires a few microliters of protein solution to be applied to an EM support grid coated with a carbon or gold film that contains numerous micrometer-scale holes. The majority of solution is then blotted away with filter paper, leaving a ~500 to 1000 Å thick aqueous film that is rapidly frozen by plunging into a bath of cryogen. Despite its enormous success, the method has several drawbacks. First, the process is remarkably wasteful. For example, with a 1 MDa protein complex that requires 3 μL of sample at 3 mg/mL for an optimal density of particles in images, the applied solution contains 9×10^−6^ g of protein consisting of 5×10^12^ molecules. If blotted to a 1000 Å thick film that covers the 3 mm diameter grid, only 7×10^−10^ g of protein or 1×10^9^ molecules remain, representing a loss of 99.98% of the material used in the experiment. At present, a typical analysis may include 10^6^ molecules, which corresponds to 0.00002 % of the sample applied. Second, the process of blotting specimens with filter paper typically takes several seconds. During this time, the surface area to volume ratio of the specimen increases dramatically, allowing protein interactions with the air-water interface that lead to preferred orientations or denaturation of the sample (Glaeser *et al.*, 2016; D’Imprima *et al.*, 2019; Noble *et al.*, 2018). The slow speed of the blotting process also complicates the design of experiments intended for time-resolved visualization of specimens.

Several recent approaches have been described to circumvent some or all of the limitations of the blotting method for specimen preparation. Time-resolved EM experiments have been performed with pressure driven microfluidic (Feng *et al.*, 2017; Kaledhonkar *et al.*, 2018) and voltage-assisted (Kontziampasis *et al.*, 2019) spraying devices. Sample loss has been addressed by using microcapillaries to apply small volumes of specimen solutions, with the *cryoWriter* device capable of applying as little as 20 nL of sample to a conventional EM grid (Kemmerling *et al.*, 2013; Arnold *et al.*, 2016, 2017; Schmidli *et al.*, 2019). This method has allowed for the impressive determination of a structure of the human 20S proteasome to ~3.5 Å resolution after isolation from 1 μL of cultured human cells (Schmidli *et al.*, 2019). Another recently described device applies a specimen to a cryo-EM grid with a purpose-constructed piezoelectric transducer designed to create an aerosol from the specimen by production of a standing wave (Ashtiani *et al.*, 2018). A highly developed approach that reduces both the issues of inefficient specimen usage and the long delays between specimen application and freezing was developed at the NIH National Resource for Automated Molecular Microscopy (Jain *et al.*, 2012; Dandey *et al.*, 2018). The *Spotiton* device uses a piezoelectric transducer coupled to a microcapillary, similar to an ink-jet printer, to apply a stream of droplets that are tens of picolitres onto an EM grid. A recent breakthrough by the same group that dramatically improves the quality of the resulting ice films was the development of ‘self-wicking’ EM grids (Wei *et al.*, 2018). These grids are produced by treating copper-rhodium EM grids with an ammonium persulfate solution at basic pH, which leads to growth of Cu(OH)_2_ nanowires on the copper surface of the grid. The nanowires function to remove excess liquid from the grid, similar to blotting paper, but at a smaller scale and faster. Unfortunately, the process of growing nanowires damages commercially available specimen grids that are already coated with holey carbon or gold films. However, plastic films can be readily produced with regular arrays of holes using a microcontact printing approach (Chester *et al.*, 2007; Marr *et al.*, 2014). These films can be deposited on the self-wicking grids and subsequently coating with either carbon or gold (Meyerson *et al.*, 2014; Russo & Passmore, 2014). The *Spotiton* device allows for rapid specimen preparation, dramatically reduces the amount of specimen required, and in principle allows for time-resolved experiments.

Several aspects of the approaches described above require precise engineering of mechanical and optical components, complicating construction in laboratories that do not specialize in instrument design. Here we present a simple and inexpensive cryo-EM specimen preparation device where samples are applied to self-wicking EM grids as droplets generated from components adapted from a readily available ultrasonic humidifier. The device has several advantages, including rapid specimen preparation, low sample consumption, and the capability of performing time resolved experiments. Most importantly, the device can be easily adapted for the design of application-specific experiments. In order to facilitate this role, the device is easily constructed from standard components and controlled by an inexpensive Raspberry Pi single board computer running a Python program. The designs for all of the custom parts are freely available as open source files and the parts can be produced by online 3D printing, CNC milling, and printed circuit board production services. This design philosophy will allow other research groups to adapt the device for other applications. Due to the use of a piezoelectric transducer for producing droplets of sample, and in homage to the *Spotiton* instrument that pioneered the use of self-wicking grids, we name our device *Shake-it-off* or *SIO*.

## Methods and Results

### Overview and process control

The *SIO* device uses four main components to prepare specimens: a piezoelectric sprayer, self-wicking grids, a plunging solenoid with detachable tweezers, and a cryogen bath. The timing of the different electronic processes is controlled by a Raspberry Pi (RPi) single board computer (raspberrypi.org) running the default Raspbian operating system. Python software on the RPi controls the voltage of the RPi’s general-purpose input output (GPIO) pins, allowing them to be set “high” (3.3 V) or “low” (0 V). A high signal from the GPIO pin applied to the gate of an N-channel enhancement mode power MOSFET (metal oxide semiconductor field effect transistor) allows a transistor to be used as a high-speed switch, with multiple transistors providing power to the different components of the device when needed (Fig. 1A). A printed circuit board (Fig. 1B) was designed with *Altium Designer* and manufactured by www.pcbway.com. A parts list summarizing all components is provided as Table 1.

**Table 1.** Components and sources for *SIO* device.

|  |
| --- |
| **Electronic components** (www.digikey.com part number): 40 position header (S9175-ND), 40 channel cable (M3AAA-4006J-ND), 12 position header (S7074-ND), 6 position header (S7039-ND), 2 position header (SAM15241-ND), 2 position shunt connector (S9341-ND), 2 mm × 5.5 mm power jack (CP-037AH-ND), [4×] pulldown resistors 10 kΩ (RNCP0603FTD10K0CT-ND), [6×] 0 Ω resistors (RMCF0603ZT0R00CT-ND), 5 V DC to DC linear regulator (1951-2747-ND), 12 V DC to DC linear regulator (1951-2739-ND), [4×] MOSFET (FQP30N06L-ND), [3×] 2 wire terminal block (A98333-ND), [2×] 3 wire terminal block (A98334-ND), [2×] diode (497-5765-1-ND), [2×] 3 pin female header connector for reed switch and infrared obstacle avoidance sensor (1568-1925-ND) |
| **Solenoids:** 24V 30 mm travel (Kuhnke: HD8286-R-F), 12V 5 mm travel (Digikey: 1528-1551-ND) |
| **Humidifier:** Universal USB Portable Mini Donut Float Water Humidifier, eBay Seller 8.52016 (19691) |
| **Power supply:** Universal 24V 6A, AC 100-240V to DC with 5mm output jack, Amazon seller Topled Light |
| **Sensors:** Keyestudio mini reed switch module (KY-021), Keyestudio Infrared Obstacle Avoidance Sensor Module (KY-032) |
| **Structure:** Thorlabs 12 × 12" optical breadboard., [3×] Hillman heavy duty corner brace (from Canadian Tire). ¼-20 × ½" Hex cap screws to connect corner braces to optical breadboard (from Canadian Tire). [4×] Retort boss, VWR. Posts are ½" diameter aluminum rods tapped for 6-32 × ½" machine screws. |
| **Adhesives (from Canadian Tire):** Armor Coat epoxy putty repair pellets (for attachment of magnets, tweezers, piezo mount to 12 V solenoid). Lepage heavy duty contact cement (for attachment of piezo to piezo mount). |
| **Control computer:** Raspberry Pi 3B+ (www.raspberrypi.org) |
| **Custom components:** Magnetic tweezer connector (3D printed in polycarbonate; www.uoftmedprint.com), piezo mount (3D printed in polycarbonate), [2×] support platform for small solenoid and IR obstacle avoidance sensor (3D printed in polycarbonate), cryogen contain (CNC milled; www.3dhubs.com), printed circuit board (www.pcbway.com). |

**Figure 1.**
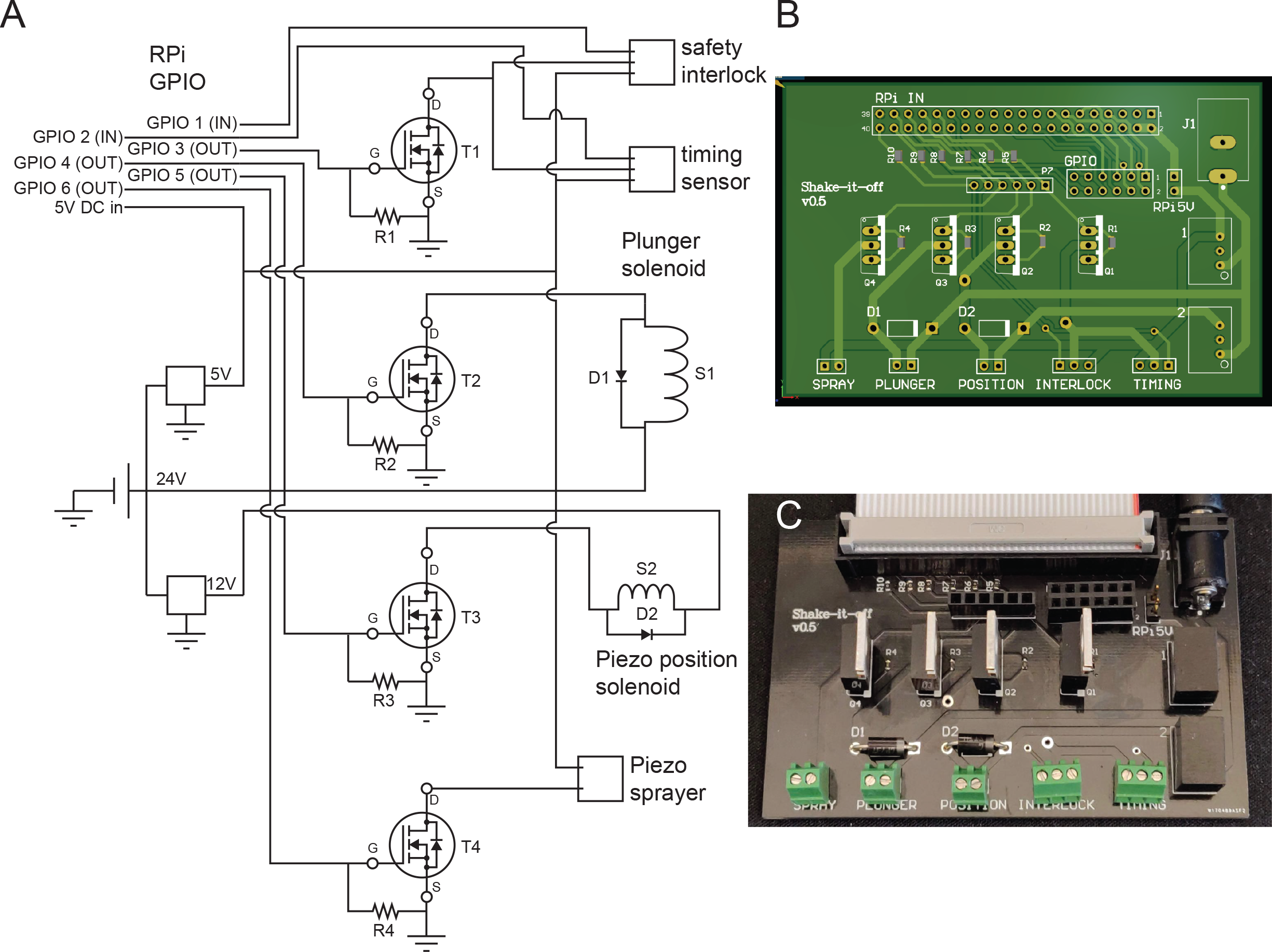
**A**, Schematic diagram of the *SIO* control circuit. **B**, Design for fabrication of custom printed circuit board. **C**, Custom fabricated printed circuit board.

### Ultrasonic dispensing of solutions

*SIO* sprays a stream of small droplets onto a self-wicking EM grid with the simple and inexpensive piezoelectric transducer and high-frequency generating circuitry of a commercially available novelty ultrasonic humidifier (Fig. 2). The humidifier was designed to be powered from the 5 V USB port of a computer. It senses when it is placed in water due to conductivity of the water connecting an electrode in its printed circuit board to ground. To override this safeguard in the humidifier circuit, a wire was added to ground the electrode (Fig. 2B, circled in blue) so that the circuit can be activated by connection to a 5 V power supply. The humidifier’s piezoelectric element is covered by a surface with numerous small holes (Fig. 2C). In order to spray a grid, the piezoelectric element is held in close proximity to the grid and protein sample (~1 μL) is applied to the surface of the transducer that is facing away from the EM grid. When the RPi activates the high-frequency generating circuit, a small stream of droplets is ejected from the piezoelectric element toward the grid. The droplet size from this specific ultrasonic humidifier has not been characterized experimentally. However, analysis of related devices found that they produce a distribution of droplets 10 to 100 nm in diameter (Kudo *et al.*, 2017) as well as a population of larger droplets 1 to 10 μm in diameter (Kudo *et al.*, 2017; Rodes *et al.*, 2007). The precise size of droplets varies slightly with the salt concentration of the solution as well as the resonant frequency of the piezoelectric transducer (Kudo *et al.*, 2017). The signal generating circuit for the piezoelectric transducer used in the *SIO* device was observed to generate a crude 108 kHz signal with a duty cycle of ~11 %. We found that a 2 μL drop was fully aerosolized after ~40 pulses of 5 msec, suggesting that each 5 msec pulse dispenses ~50 nL. Application of ~1 μL to the transducer is necessary to cover the array of small holes on the transducer used to generate droplets. In principle, the ~1 μL of applied sample can be used to prepare numerous specimen grids.

**Figure 2.**
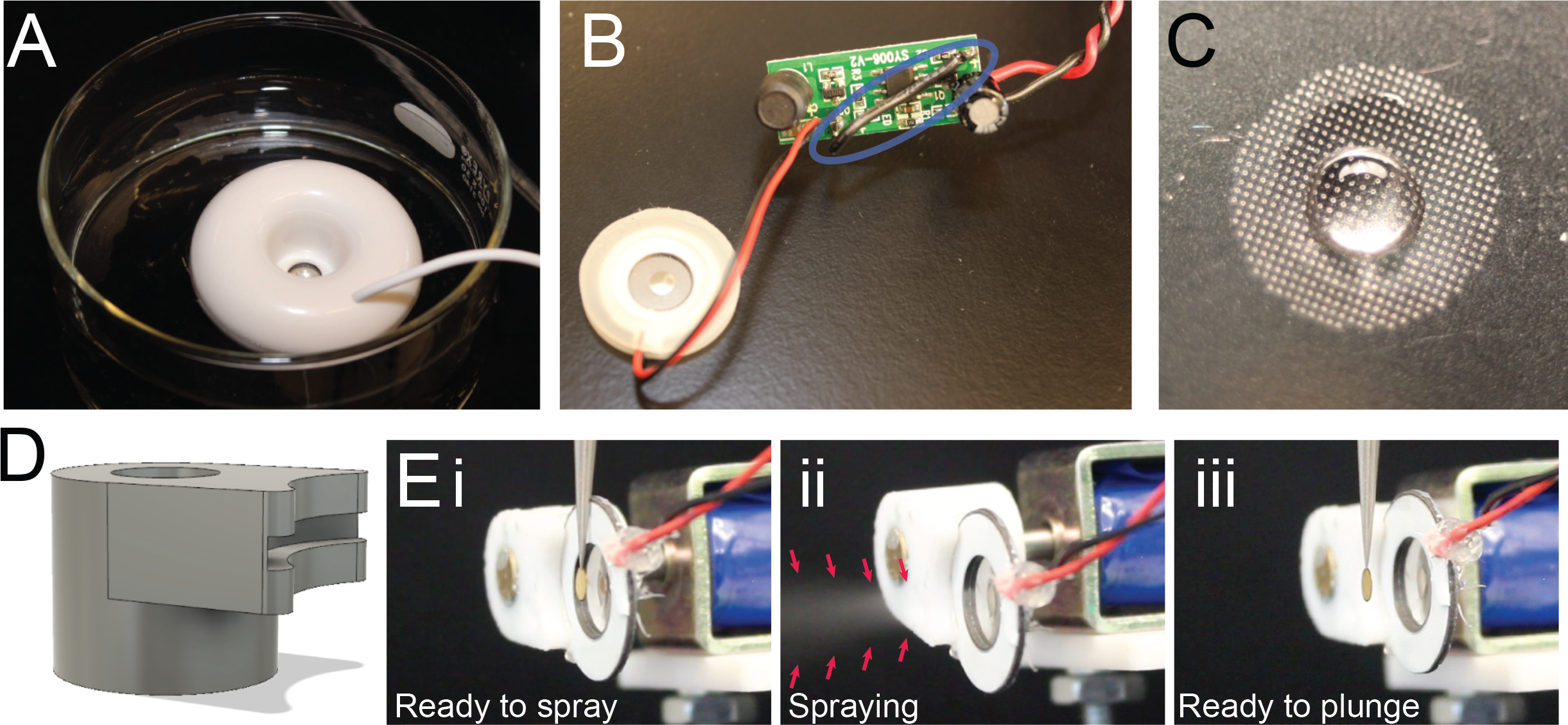
**A**, Novelty USB ultrasonic humidifier containing a piezoelectric element and high-frequency generating circuit used to spray specimen. **B**, The piezoelectric transducer and high-frequency generating circuit extracted from the humidifier shown in part A. A wire that allows the circuit to be activated out of water is circled in blue. **C**, A droplet (1 μL) applied to the piezoelectric element surface opposite to direction of liquid spray. **D**, 3D design for piezoelectric element mounting connector, printed in polycarbonate. **E**, Piezoelectric element, tweezers, and grid in the specimen application position (**i**), showing spray from the piezoelectric transducer (**ii**), and with the piezoelectric element retracted in the ready-to-plunge position (**iii**).

### Self-wicking grids

Self-wicking grids were prepared as described previously by incubating 400 mesh Cu/Rh grids (Maxtaform, Electron Microscopy Sciences) in nanowire growth solution consisting of 750 mM NaOH and 65 mM (NH_4_)_2_S_2_O_8_ for 5 min followed by washing with water (Razinkov *et al.*, 2016; Wei *et al.*, 2018). A continuous formvar film with ~2 μm holes spaced 4 μm centre-to-centre was prepared and overlaid on the self-wicking grids as described previously (Marr *et al.*, 2014), and coated with ~35 nm of gold with a Leica EM ACE200 sputter coater before dissolving the plastic film with chloroform. It may be that during spraying of the grid, droplets smaller than the hole size in the gold film pass through the holes, while the larger droplets cover the holes. However, the high density of smaller droplets may also contribute to the formation of a suitable film for cryo-EM once holes are blocked by larger droplets.

### Grid plunging

Reproducibly hitting the EM grid with the stream of droplets proved challenging when the grid was held sufficiently far from the piezoelectric transducer to allow plunging of the tweezers. Therefore, we mounted the piezoelectric element on a small “pull type” 12 V solenoid. The mount attaching the piezoelectric element to the solenoid was designed with *Autodesk Fusion 360* software (Fig. 2D). All 3D printed components were printed in polycarbonate with a Fortus 450mc 3D printer at the University of Toronto Medstore 3D printing facility (www.uoftmedprint.com). The solenoid is positioned so that in the spraying position the piezoelectric element touches the tweezers and is ~1 mm away from the grid (Fig. 2E, part i), allowing the spray from the piezoelectric element to hit the grid (Fig. 2E, part ii). Removing power from the solenoid allows the spring-driven retraction of the piezoelectric element from the grid (Fig. 2E, part iii), providing the necessary clearance for the grid to be plunged into cryogen without the tweezers hitting the element. Removing current from an inductive load such as a solenoid can produce a sudden voltage spike across the load. In order to avoid damage to the MOSFET from this voltage spike, a “fly-back” diode was included in the circuit with a polarity opposite to the applied voltage (Fig. 1A, Diodes 1 and 2).

Plunging of the grid is performed by a large solenoid with a 30 mm travel distance, identical to the solenoid previously used for the same purpose in the *cryoWriter* device (Arnold *et al.*, 2016; Kemmerling *et al.*, 2013; Arnold *et al.*, 2017). A fly-back diode was also employed for this solenoid. In order to allow fast and convenient connection and disconnection of tweezers from the plunging solenoid, a 3D printed magnetic connector was designed (Fig. 3A). This connector has one half attached to the plunging solenoid by a hexagonal nut held in place with epoxy putty, while the other half is attached to the blunt end of the tweezers, also by epoxy putty. The two halves are reversibly attached together by 6 mm diameter 3 mm thick cylindrical neodymium magnets epoxied into each half.

**Figure 3.**
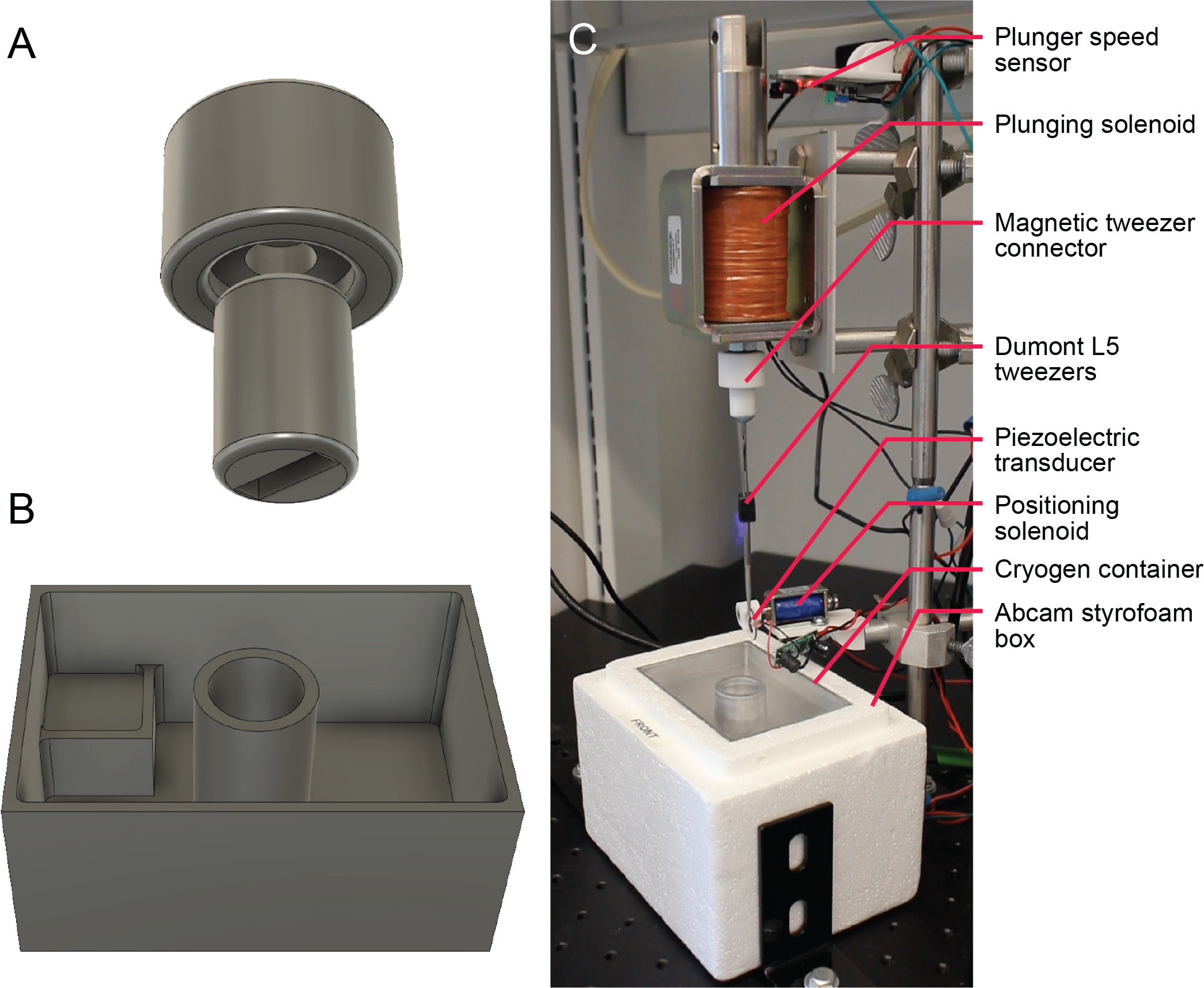
**A**, 3D design for magnetic tweezer connector. The two parts are held together by embedded magnets and the components are printed in polycarbonate. **B**, 3D design for the 99×70×43 mm cryogen container, which fits within a standard Abcam Styrofoam box, milled from 6061 alloy aluminum. **C**, Fully assembled *SIO* specimen preparation device.

### Cryogen container

Freezing of specimens for cryo-EM is typically done by rapid plunging into liquid ethane cooled by liquid nitrogen. Ethane freezes at liquid nitrogen temperature and consequently cryogen containers for specimen preparation devices need to be designed to keep the ethane in its liquid state. One method for keeping ethane liquid is immersion of a heating element, as done in many devices (Bellare *et al.*, 1988). Alternatively, the cryogen container can be carefully designed so that the thermal contact between the ethane chamber and liquid nitrogen chamber is sufficient to keep the ethane liquid, but not sufficient to freeze the ethane, as done with the popular *Vitrobot* grid freezing device. This approach removes the need for an immersed heating element but prevents the cryogen container from being manufactured from materials with high thermal conductivity such as aluminum. Tivol and colleagues (Tivol *et al.*, 2008) presented an ingenious solution to this problem: mixing propane with ethane in an ~60:40 ratio depresses the freezing point of both alkanes so that the mixture does not freeze even at liquid nitrogen temperature. We designed a liquid nitrogen reservoir with a propane/ethane bath and manufactured it from a single block of 6061 alloy aluminum by computer numerical control (CNC) milling (www.3dhubs.com). The cryogen container (Fig. 3B) was designed so that it could be insulated by a small Styrofoam box used as a standard shipping container by the company Abcam. These containers are easily found in many biological research laboratories. In the complete *SIO* device, the cryogen container is held in place relative to the piezoelectric transducer and plunging solenoid by corner braces on an optical bread board (Thorlabs) using a vertical post and standard retort stand boss heads (Fig. 3C).

Accidentally plunging the sharp tweezers before the cryogen container is in place presents a safety hazard. Therefore, we added a safety interlock to the design. A neodymium magnet was embedded in the Styrofoam of the cryogen container and a miniature reed switch attached to the positioning corner brace for the container. The magnet is held near the switch only when the container is positioned appropriately. The field from the magnet closes the switch, which is detected by the control software on the RPi. The software only allows the plunger solenoid to be activated if the cryogen container is in place.

## Structure determination

Conditions were identified in which ice suitable for cryo-EM structure determination could be produced consistently using horse spleen apoferritin (Sigma). The full specimen preparation process is shown in Video 1. The 50 % glycerol in the sample was reduced by diluting 20× in TBS (50 mM Tris-HCl, 150 mM NaCl, pH 7.4) and concentrating 20× with a centrifuged concentrating device with a 100 kDa molecular weight cut-off membrane. Immediately before use, grids were glow discharged in air for 2 min at 40 mbar with 25 mA current in a Pelco easiGlow glow discharge device, with the nanowire surface of the grid facing the plasma of the discharge device. A 1 μL droplet of apoferritin solution at 5 mg/mL in TBS was placed on the back of the piezoelectric transducer. To apply specimen to the grid, the high-frequency generating circuit was energized for 5 msec. At 5 msec the piezo was retracted from the grid by deenergizing the piezo positioning solenoid and the plunging solenoid was energized. An infrared obstacle avoidance sensor (keyestudio) mounted on the plunging solenoid, as well as recording of a slow-motion video at 2.5 msec/frame with a mobile phone video camera (OnePlus 6), both suggested a total time from the beginning of spraying to complete immersion of the grid in cryogen of ~90 msec (Video 2).

Grids were characterized with a FEI Tecnai F20 microscope operating at 200 kV and micrographs were recorded as 15 sec movies of 30 frames at 5 electrons/pixel/sec and a calibrated pixel size of 1.45 Å/pixel. A dataset of 324 high-resolution movies was collected on a Titan Krios G3 electron microscope operating at 300 kV and recorded as 180 frames over 30 s on a Falcon 3EC with an exposure rate of 0.786 electrons/pixel/sec, a calibrated pixel size of
1.06 Å/pixel, and a total exposure of 21 electrons/Å^2^. Atlas images of grids (Fig. 4A and B) show that the device creates an area of thick ice surrounded by a band of grid squares that have ice suitable for data collection. An image of a grid square (Fig. 4B, inset) shows that edges of the grid square can have holes without ice, possibly due to excess material being removed by the self-wicking grids, while the centre of the grid square has ice suitable for imaging. Apoferritin particles can be seen in images from holes in this region (Fig. 4C). Image analysis with *cryoSPARC* (Punjani *et al.*, 2017) allowed selection of 308,752 particle images that were reduced to a dataset of 172,469 particle images by 2D and 3D classification. These images were used to calculate a 3D map at 2.6 Å resolution map (Fig. 4D).

**Figure 4.**
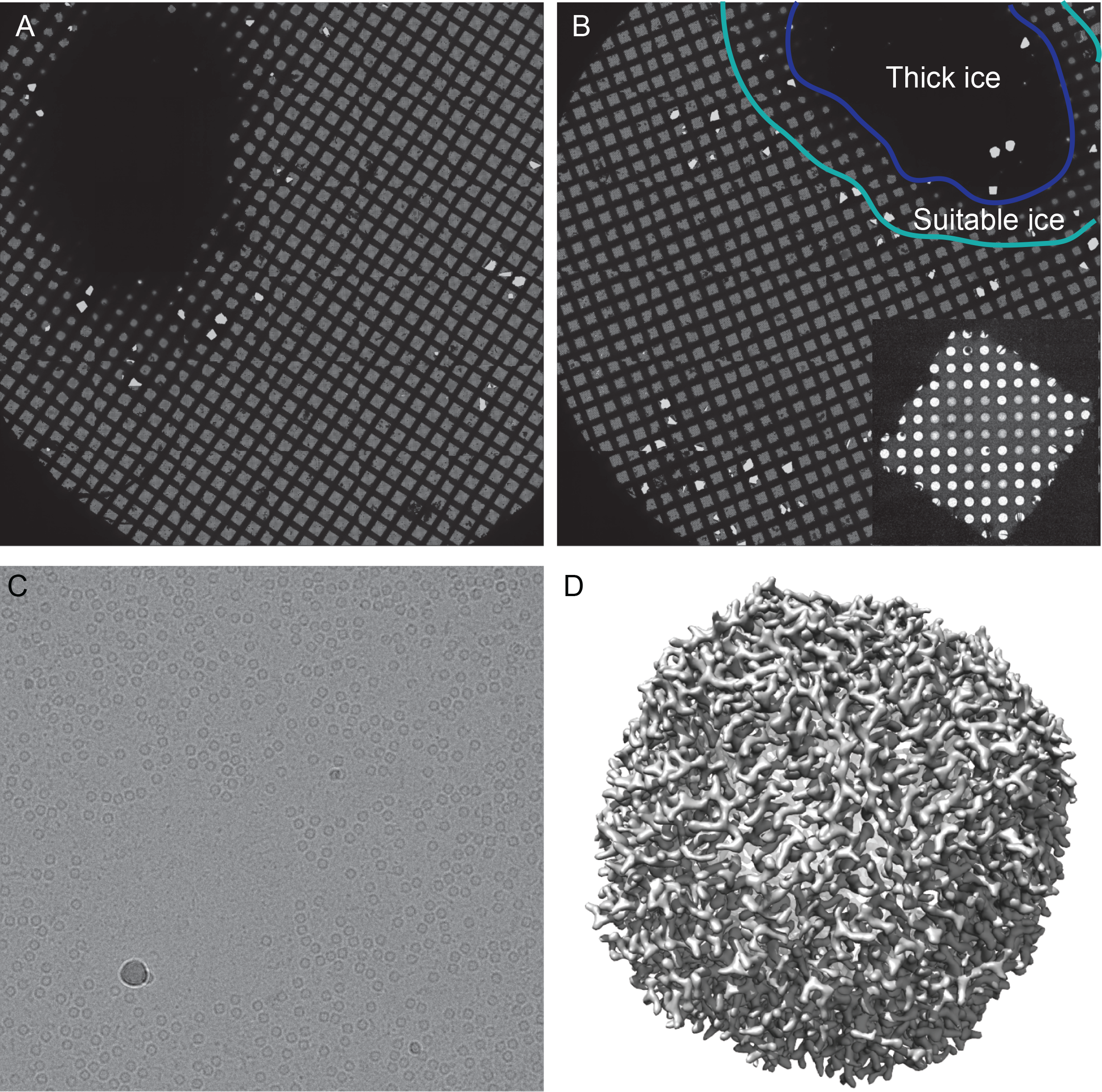
**A and B**, Grid atlases from the Titan Krios microscope showing a large area of thick ice (circled in purple in part B) with a peripheral area of ice suitable for data collection (circled in cyan in part B). Inset: a grid square from a region with suitable ice for imaging showing some over-wicking at its periphery. **C**, An image from the TF20 microscope showing equine apoferritin particles in ice. **D**, A 3D map of equine apoferritin at 2.6 Å resolution from the Titan Krios microscope.

## Discussion

Here we have shown that a simple device combining 3D printed and CNC-milled parts, a printed circuit board, and inexpensive consumer components, is capable of producing specimens suitable for high-resolution cryo-EM analysis with ~100 msec time resolution. This time resolution can allow for simple mix-and-freeze experiment by, for example, placing two separate ~500 nL drops of reactant on the piezoelectric transducer, rather than a single ~1 μL drop. All designs for the 3D printed and CNC-milled parts of the device, as well as the design files for the printed circuit board and the Python instrument control software, are available at https://github.com/johnrubinstein. With these designs, scientists with limited access to technical workshops and equipment can easily construct their own device by making use of services accessed over the internet.

The intention of this open source design is that the *SIO* device can be easily customized and modified for specialized applications. Various components of the device may also warrant improvements. For example, a second version of the device is already planned to allow easier positioning of the piezoelectric transducer relative to the self-wicking grid by making use of a 3D printed *xyz*-positioning stage. Further, a simple blotting version of the device is also being constructed that will provide the advantages of commercially available grid freezing devices such as the *Vitrobot*, Gatan *CP3*, and Leica *EM GP* at a fraction of the cost and with various desired customizations.

## Contributions

JLR conceived the project, design and built most of the device, wrote the software, tested the device, and wrote the manuscript. HG imaged the majority of the specimens by cryo-EM, made numerous suggestions for the improvement of the device, and calculated 3D maps. ZAR tested the device, suggested improvements, and calculated 3D maps. AH suggested improvements to the electronics and designed the printed circuit board. AA and CMY suggested aspects of the design, made components used for optical characterization of the spraying process that are not described here, and guided initial design of electronics. JMDT re-designed some of the 3D printed components. SB assisted with Titan Krios data collection. TK provided suggestions on design and performed several workshop steps.

## Acknowledgements

We thank Alex Wei, Venkat Dandey, Clint Potter, and Bridget Carragher for helpful conversations related to self-wicking grids and the *Spotiton* device, as well as Luca Rima and Thomas Braun for helpful conversations related to the *cryoWriter* device. JLR thanks Jianhua Zhao, Anna Zhou, and Daniel Schep for the gift of a Raspberry Pi computer that started this project.

---

**Video 1.** Illustration of the full specimen preparation process.

**Video 2.** Video at 480 frames per second showing a grid being plunged into liquid.

